# Interleukin-1 regulates follicular T cells during the germinal center reaction

**DOI:** 10.1101/2023.03.29.534687

**Authors:** Aude Belbezier, Paul Engeroff, Gwladys Fourcade, Helene Vantomme, Romain Vaineau, Bruno Gouritin, Bertrand Bellier, Isabelle Brocheriou, Nicolas Tchitchek, Stephanie Graff-Dubois, David Klatzmann

**Affiliations:** Sorbonne Université, INSERM, Immunology-Immunopathology-Immunotherapy (i3), F-75005, Paris, France; AP-HP, Hôpital Pitié-Salpêtrière, Biotherapy (CIC-BTi) and Inflammation-Immunopathology-Biotherapy Department (i2B), F-75651, Paris, France; AP-HP, Hôpital Pitié-Salpêtrière, Department of Pathology, F-75651, Paris, France

**Author notes:** co-senior authors (Stephanie Graff-Dubois, David Klatzmann).

## Abstract

Antibody production by plasma cells and generation of long-term memory B-cells are positively regulated by Tfh cells and negatively regulated by Tfr cells in germinal centers. However, the precise role of Tfr cells in controlling antibody production is still unclear. We have previously shown that Tfh and Tfr cells both express IL-1R1, while only Tfr cells express the IL-1R2 decoy and IL-1Ra antagonist receptors. To study the role of these receptors in the regulation of the B cell response by Tfh and/or Tfr cells, we generated mice with knockout of IL-1 receptors in Tfh and/or Tfr cells and measured antibody production and cell activation upon immunization. We showed that IL-1β concentration is increased in the draining lymph node after immunization. Antigen-specific antibody levels and cell activation phenotypes indicated that IL-1β can activate both Tfh and Tfr cells through IL-1R1 stimulation. Surprisingly, IL-1R2 and IL-1Ra expression on Tfr cells does not block IL-1 activation of Tfh cells, but rather prevents IL-1/IL-1R1-mediated early activation of Tfr cells. IL-1Rs similarly regulate antibody response against autoantigens and its related pathophysiology in an experimental lupus model. Altogether, these results show that IL-1R1 inhibitory receptors expressed by Tfr cells prevent their own activation and suppressive function, thus licensing IL-1-mediated activation of Tfh cells after immunization.

**One Sentence Summary:** Functional knockout of IL-1 agonist (R1) and antagonist (R2 and Ra) receptors reveals that IL-1-R2 and -Ra on Tfr cells prevent their own early activation to license the expansion and activation of Tfh cells after immunization.

## Introduction

Interleukin-1α and β (IL-1), the two first discovered members of the IL-1 family, are mainly known as potent proinflammatory cytokines in innate immunity (Chan and Schroder, 2019; Dinarello, 2011; Garlanda et al., 2013). IL-1α is predominantly produced by pyroptotic cells and cells undergoing specific inflammasome-independent forms of cell death (Cohen et al., 2010), while IL-1β is produced by hematopoietic cells such as blood monocytes and tissue macrophages in inflammatory conditions (Garlanda et al., 2013). All functions triggered by IL-1α and β are mediated through the agonist IL-1 receptor IL-1R1 (Dower et al., 1985). This interaction is tightly regulated by two other IL-1 receptors: (i) IL-1Ra, a soluble IL-1 receptor antagonist that binds to IL-1R1 and prevents the binding of IL-1 α/β and (ii) IL-1R2, a decoy of the IL-1 receptor located mainly in the membrane, which, when binding to IL-1, does not induce signal transduction, unlike the IL-1R1 receptor, thus reducing IL-1 availability (Garlanda et al., 2013). IL-1α and β are mainly considered as key triggers of immediate inflammation that have common pleiotropic functions, such as triggering fever, hematopoiesis or leukocyte attraction (Chan and Schroder, 2019). However, recent work has suggested that they could also be implicated in the T follicular helper cell (Tfh) and T follicular regulatory cell (Tfr) regulation of B cell immune responses and thus in the adaptive immune response (Ritvo et al., 2017; Ritvo and Klatzmann, 2019).

Activated B cells undergo mutation, selection and affinity maturation in germinal centers (GCs), which leads to antibody (Ab) production by plasma cells and the generation of long-term memory B-cells. B-cell activation can occur independently of T cells or in T-cell-dependent fashion within GCs (De Silva and Klein, 2015). GCs are positively regulated by Tfh and negatively regulated by Tfr (Vinuesa et al., 2016; Sage et al., 2016). Tfh cell-derived CD40L expression, IL-4 and IL-21 play essential roles in GC B cell proliferation, survival and affinity maturation (Vinuesa et al., 2016). The origin and mode of action of Tfr cells are more controversial. They are thought to derive from regulatory T cells (Tregs) and to act by regulating the help Tfh cells provide to B-cells (Sage and Sharpe, 2016; Maceiras et al., 2017; Wollenberg et al., 2011; Linterman et al., 2023).

We recently showed that Tfh cells express the IL-1R1 agonist receptor while Tfr cells also express IL-1R1, albeit at lower levels than Tfh cells, but also high levels of the IL-1R2 decoy receptor and IL-1Ra, the IL-1 receptor antagonist (Ritvo et al., 2017). Furthermore, a role of an IL-1/IL-1Rs axis in B cell responses is supported by the fact that antibody production is (i) reduced in IL-1–deficient mice (Nakae et al., 2001a; b) and (ii) enhanced by adding IL-1 during immunization (Reed et al., 1989; Staruch and Wood, 1983; Nencioni et al., 1987) or in mice lacking the expression of IL-1Ra (Nakae et al., 2001a; b).

These observations raised questions about the specific role of each IL-1 receptor in Tfh and Tfr cell balance and, consequently in antibody production. To address these questions, we generated mice specifically lacking IL-1 receptors in Tfh and/or Tfr cells. We showed that IL-1 activates Tfh and Tfr cells through IL-1R1, whereas IL-1Ra and IL-1R2 expressed on Tfr cells inhibit their own activation. These results demonstrate a major role for IL-1 not only in the positive regulation, but also in the negative regulation of humoral immunity.

## Results and discussion

### IL-1 mediates Tfh and Tfr cell activation

In CD4^+^ T cells, IL-1R1 is mostly expressed by Tfh cells and at a lower level on Tfr cells (Nakae et al., 2001). We crossed CD4^cre^ mice with IL-1R1^lox^ mice and observed a proper deletion of IL-1R1 on CD4^+^ cells using reverse transcription-polymerase chain reaction (RT-PCR) (Fig. S1A).

At steady state, CD4^cre^IL-1R1^lox^ mice had normal numbers of Tfh, Tfr or Treg cells, and normal numbers of transitional B cells (CD19^+^CD38^-/low^B220^low^IgD^-^GL7^-^Bcl6^-^), GC B cells (GCBs) (CD19^+^GL7^+^Bcl6^+^IgD^-^) or plasmablasts (CD19^+^CD38^+^B220^low^IgG^+^IgD^-^). However, they had significantly reduced serum IgG Ab levels at steady state (Fig. 1A). When CD4^cre^IL-1R1^lox^ mice were immunized with ovalbumin (OVA), four weeks after immunization we observed a significant reduction of the proportion of Tfh but not of Tfr cells, leading to a significantly increased Tfr/Tfh ratio in comparison with control mice (Fig. 1B). This was accompanied by a markedly decreased production of GCBs (Fig. 1B). These effects translated into a significant decrease of anti-OVA Ab production (Fig. 1C), whereas the ratio of OVA-specific Ab to total Ab was not affected (Fig. 1D). These results indicate an effect of IL-1R1 on the Tfh/Tfr control of humoral responses.

**Fig. 1.**
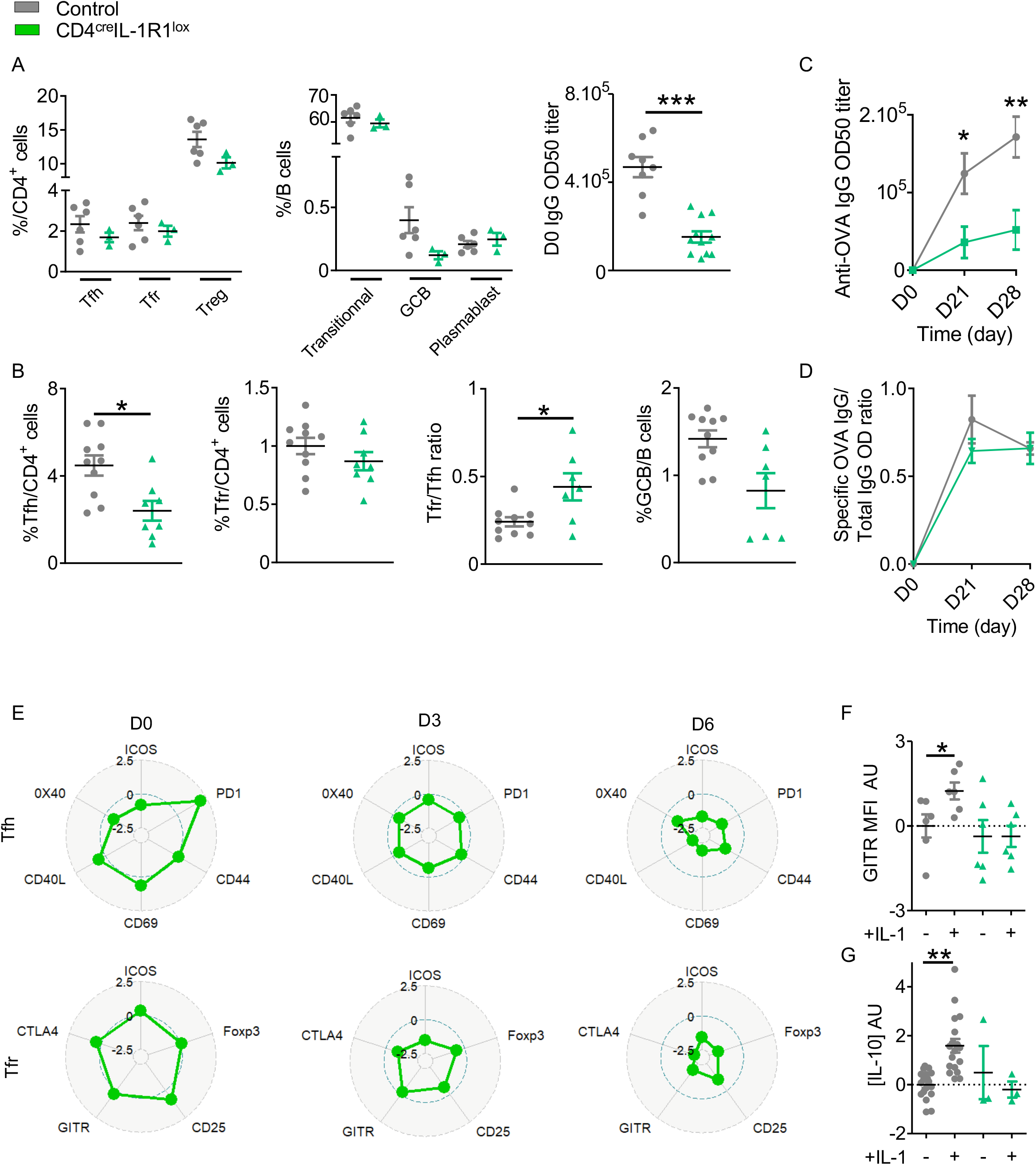
IL-1 regulates Tfh cell but also Tfr cell activation. (A) Quantification of CD4^+^ T and B cell populations within SLOs and circulating IgG antibody levels before any immunization in CD4^cre^IL-1R1^lox^ (green triangle) and CD4^cre^ (grey circle) mice. (B) Proportion of Tfh, Tfr, GCB cells and Tfr/Tfh ratio 28 days after intraperitoneal immunization with OVA-Alum in CD4^cre^IL-1R1^lox^ (green triangle) and CD4^cre^ mice (grey circle). (C) Circulating anti-OVA IgG antibody level after intraperitoneal immunization with OVA-Alum in CD4^cre^IL-1R1^lox^ (green triangle) and CD4^cre^ (grey circle) mice from D0 to D28. (D) Ratio of specific anti-OVA IgG Ab/total IgG Ab after OVA immunization in CD4^cre^IL-1R1^lox^ (green triangle) and CD4^cre^ (grey circle) mice from D0 to D28. (E) Tfh and Tfr expression levels of the indicated markers in Foxp3^GFP^CD4^cre^IL-1R1^lox^ mice (green) compared to their expression in cells from Foxp3^GFP^ mice after immunization at the indicated time point. Expression levels are indicated in arbitrary units that correspond to data centered and reduced relative to cells from Foxp3^GFP^ mice. (F, G) Histograms comparing the MFI of GITR (F) and the level of IL-10 production (G) of Tfr cells from Foxp3^GFP^CD4^cre^IL-1R1^lox^ (green triangle) and Foxp3^GFP^ (grey circle) mice cultured with or without IL-1. GITR MFI and IL-10 levels are expressed in arbitrary units that correspond to data centered and reduced relative to Tfr from Foxp3^GFP^ (grey) mice grown without IL-1. *P < 0.05, **P < 0.01, ***P < 0.005, Mann-Whitney U test for unpaired data and Wilcoxon paired test for paired data. (B,C,D,F,G) Data are representative of three independent experiments.

To further explore the role of IL-1R1 in Tfh and Tfr cell activation, we performed quantitative and qualitative analysis of Tfh and Tfr cell populations from draining lymph nodes of OVA-immunized mice by spectral cytometry. To better discriminate the effect of IL-1R1 deletion on Tfr from Tfh cells, we generated Foxp3^GFP^CD4^cre^IL-1R1^lox^ mice. According to the literature reporting that Tfh cell expansion begins 3 days after immunization and reaches a peak after 6-7 days (Ritvo et al., 2017; Ritvo and Klatzmann, 2019; Rasheed et al., 2006), we focused our analysis on day-3 (D3) and −6 (D6) post-immunization. Before immunization, the absence of IL-1R1 expression led only to a non-significant increase of PD1 on Tfh (p= 0.1). Compared to Tfh from immunized control mice, Tfh from Foxp3^GFP^CD4^cre^IL-1R1^lox^ immunized mice showed a small decrease of activation marker expression at D3 and a marked decrease at D6. In contrast, there was a marked decrease of the Tfr activation profile already at D3, which was even more pronounced at D6 (Fig. 1E). These results indicate a role of IL-1R1 in the IL-1-mediated activation of both Tfh and Tfr cells, despite low IL-1R1 expression on the latter. Actually, we confirmed the functionality of IL-1R1 on Tfr cells in vitro. The addition of IL-1β to purified Tfr from OVA-immunized mice significantly increased IL-10 production and GITR expression (Fig. 1F, G), and such activation could no longer be detected with IL-1R1-deleted Tfr (Fig. 1F, G). Collectively, these results suggest a functional role of IL-1R1 on Tfr cells, which calls for study of the impact of specific depletion of IL-1R1 on Tfr cells.

We thus crossed Foxp3^creYFP^ mice, which express yellow fluorescent protein (YFP) as a reporter gene for the expression of Foxp3, with IL-1R1^lox^ mice. We confirmed that Foxp3^creYFP^IL-1R1^lox^ mice had a deletion of IL-1R1 expression on Tfr but not on Tfh cells using RT-PCR (Fig. S1B). At steady state, compared to control mice, Foxp3^creYFP^IL-1R1^lox^ mice presented (i) no significant modifications of Tfh, Tfr or Treg cells, nor of the B cell subsets (Fig. 2A) and (ii) a significantly lower level of circulating IgG Ab (Fig.2A). Four weeks after immunization, compared to controls, they had a significantly decreased Tfr/Tfh ratio (Fig. 2B) and an increase of specific anti-OVA Ab (Fig. 2C), with a significantly increased ratio of specific to total Ab (Fig. 2D). Altogether, these results indicate that the expression of IL-1R1 on Tfr cells is involved in their own activation.

**Fig. 2.**
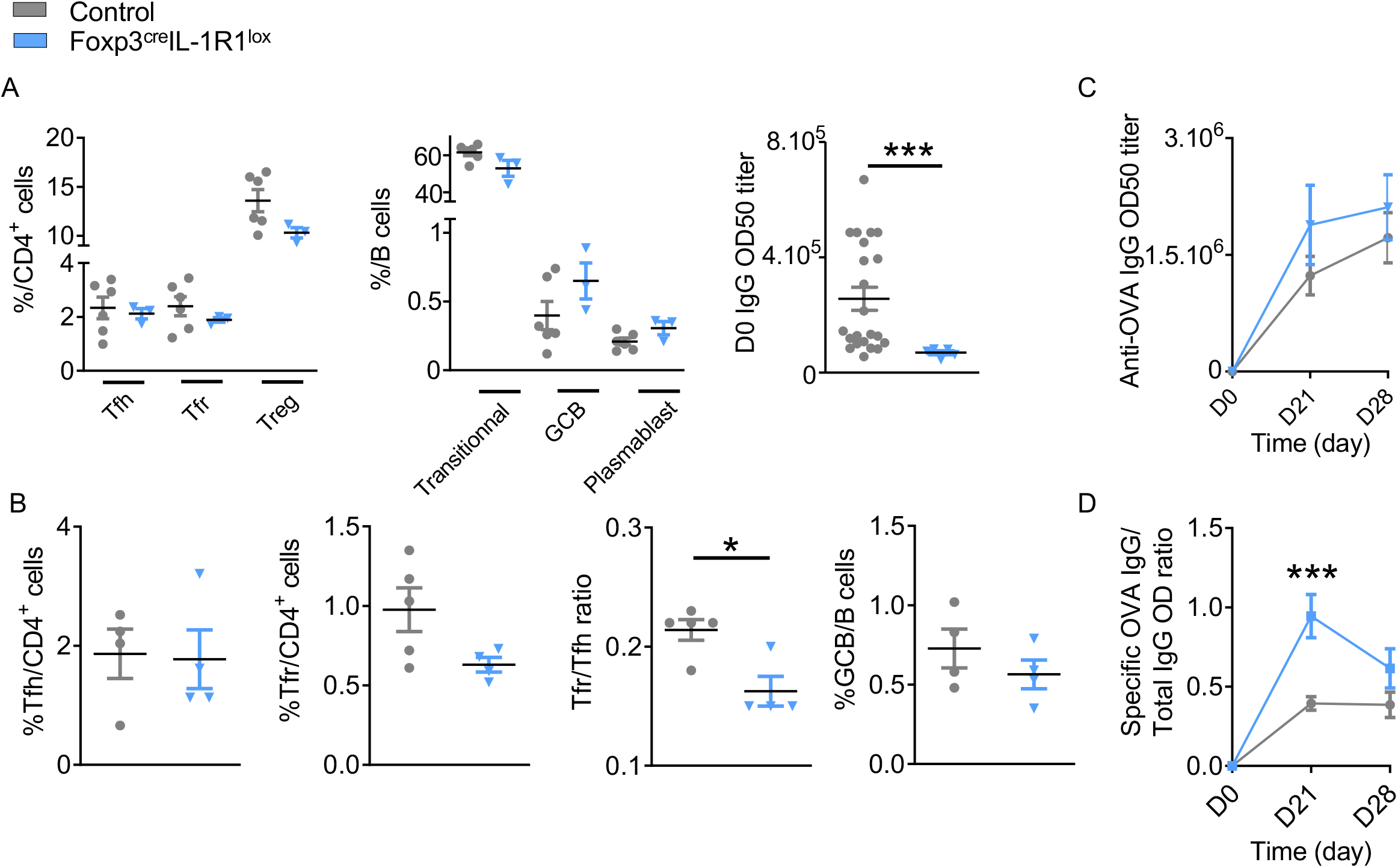
IL-1R1 receptor expressed by Tfr cells is functional and critical for the regulation of the humoral immune response. (A) Quantification of CD4^+^ T and B cell populations within SLOs and circulating IgG antibody levels before any immunization in Foxp3^creYFP^IL-1R1^lox^ (blue triangle) and Foxp3^creYFP^ mice (grey circle). (B) Proportion of Tfh, Tfr, GCB cells and Tfr/Tfh ratio 28 days after intraperitoneal immunization with OVA-Alum in Foxp3^creYFP^IL-1R1^lox^ (blue triangle) and Foxp3^creYFP^ mice (grey circle). (C) Circulating anti-OVA IgG antibody level after intraperitoneal immunization with OVA-Alum in Foxp3^creYFP^IL-1R1^lox^ (blue triangle) and Foxp3^creYFP^ mice (grey circle) from D0 to D28. (D) Ratio of specific anti-OVA IgG Ab/total IgG Ab after OVA immunization in Foxp3^creYFP^IL-1R1^lox^ (blue triangle) and Foxp3^creYFP^ mice (grey circle) at the times indicated.*P < 0.05, **P < 0.01, ***P < 0.005, Mann-Whitney U test. (B, C, D) Data are representative of three independent experiments.

The function of IL-1R on Tfh cannot be addressed specifically, but can be inferred by comparing the phenotypes of CD4^cre^IL-1R1^lox^ mice and Foxp3^creYFP^IL-1R1^lox^ mice. The absence of IL-1R1 on Tfh and Tfr prevents Tfh expansion and decreases the expression of activation markers post-immunization. However, the lack of IL-1R1 on Tfr should reduce their activation and suppressive function on Tfh, and thus cannot be responsible for the reduced Tfh activation. Thus, according to the results obtained in these mice it can be concluded that the IL-1/IL-1R1 interaction is essential to Tfh activation and expansion triggered by immunization. Indeed, loss of IL-1R1 on CD4 T cells results in defective production of anti-OVA IgG after immunization as well as defective expression of costimulatory molecules on both Tfh and Tfr cells. Moreover, at steady state, the absence of IL-1R1 on CD4 T cells translates into a reduction of the IgG titer, suggesting that IL-1R1 is required for IgG production. These data are in agreement with a complementary study in allergy where we show that IL-1/IL-1R1 signaling is necessary for the redirection of the pathogenic IgE response to the IgG response (Engeroff et al., manuscript submitted; (bior_X_iv)). Finally, while we previously overlooked the low IL-1R1 expression by Tfr, we show here that this receptor is functional. Indeed, mice lacking IL-1R1 on Tfr cells have (i) Tfrs showing a lower activation profile and (ii) a higher production of specific antibodies after immunization. Therefore, IL-1 could result in the activation of both Tfh and Tfr, which would have opposite functions.

### IL-1R2 and IL-1Ra control humoral immunity

To investigate the role of IL-1R2 in Tfr cell activation, we crossed Foxp3^creYFP^ mice with IL-1R2^lox^ mice. At steady state, we observed a non-significant increase of follicular T cells including both Tfh and Tfr in Foxp3^creYFP^IL-1R2^lox^ mice compared to control mice (Fig. 3A). However, Foxp3^creYFP^IL-1R2^lox^ mice showed a significant decrease in the proportion of transitional B cells, with a significant increase in the proportion of GCBs and plasmablasts (Fig. 3A). This suggests that deletion of IL-1R2 on Tfr specifically impacts B cell maturation, even though circulating IgGs were not affected in naïve mice (Fig. 3C). Four weeks after OVA-immunization, we observed a significant increase of the Tfr/Tfh ratio in FoxP3^creYFP^IL-1R2^lox^ mice compared to control mice, whereas the GCB/B ratio remained similar (Fig. 3B). Interestingly, FoxP3^creYFP^IL-1R2^lox^ mice produced greatly reduced levels of OVA-specific IgG compared to control mice (Fig. 3C).

**Fig. 3.**
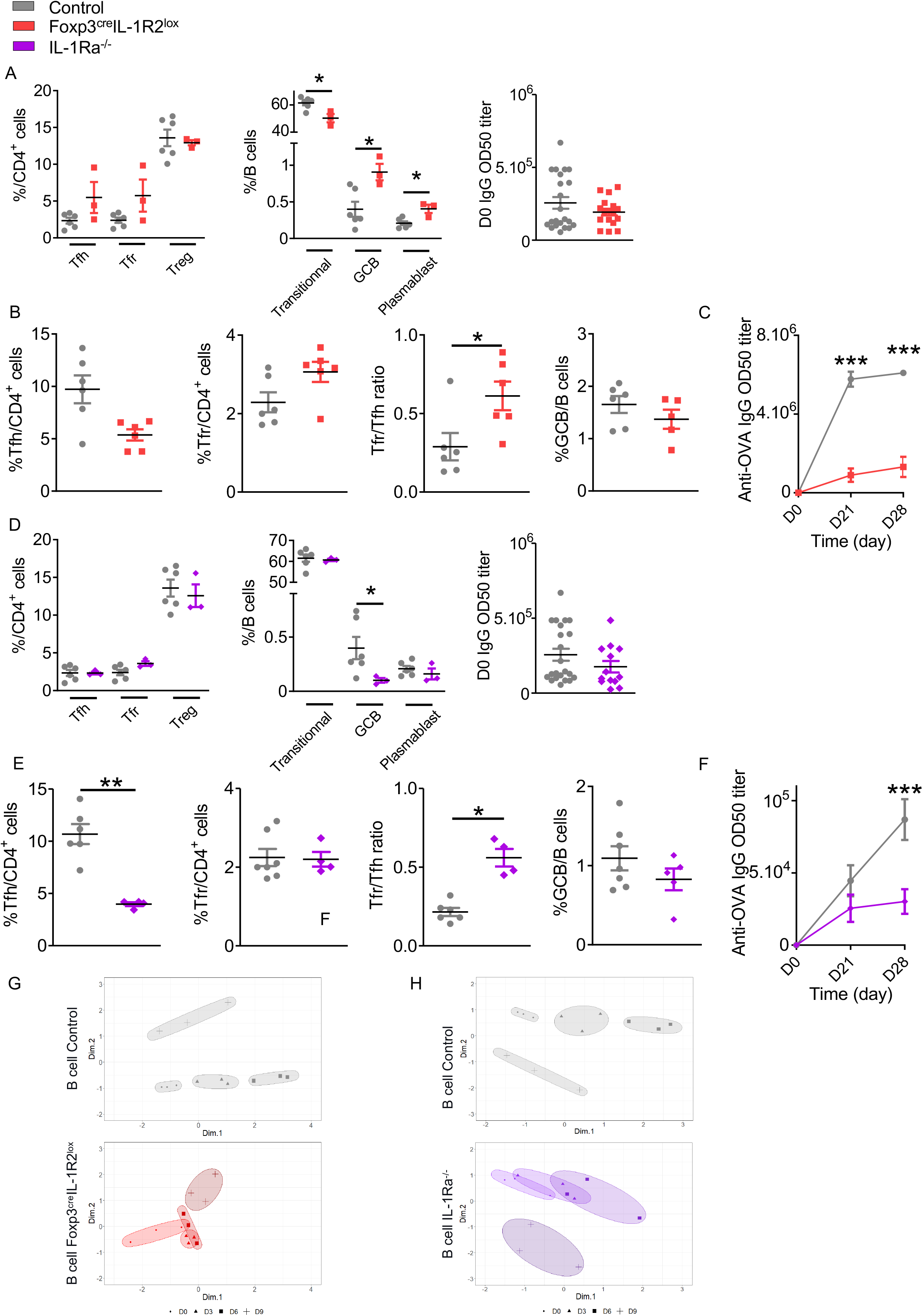
IL-1R2 and IL-1Ra receptor are functional and critical for the regulation of the humoral immune response. (A) Quantification of CD4^+^ T and B cell populations within SLOs and circulating IgG antibody levels before any immunization in Foxp3^creYFP^IL-1R2^lox^ (red square) and Foxp3^creYFP^ (grey circle) mice. (B) Proportion of Tfh, Tfr, GCB cells and Tfr/Tfh ratio 28 days after intraperitoneal immunization with OVA-Alum and (C) circulating anti-OVA IgG antibody level after intraperitoneal immunization with OVA-Alum in Foxp3^creYFP^IL-1R2^lox^ (red square) and Foxp3^creYFP^ (grey circle) mice from D0 to D28. (D) Quantification of CD4^+^ T, B cell subsets within SLOs and circulating IgG antibody levels before any immunization in Foxp3^GFP^IL-1Ra^-/-^ (purple diamond) and Foxp3^GFP^IL-1ra^WT^ (grey circle) mice. (E) Proportion of Tfh, Tfr, GCB cells and Tfr/Tfh ratio 28 days after intraperitoneal immunization with OVA-Alum and (F) circulating anti-OVA IgG antibody level after intraperitoneal immunization with OVA-Alum in IL-1Ra^-/-^ (purple diamond) and Foxp3^GFP^ (grey circle) mice. (G) MDS of B cell subset frequencies on Foxp3^creYFP^IL-1R2^lox^ (red) and Foxp3^creYFP^ (grey) mice from D0 to D28 after immunization. (H) MDS of B cell population frequencies on Foxp3^GFP^IL-1Ra^-/-^ (purple) and Foxp3^GFP^ (grey) mice at the indicated time point after immunization. *P < 0.05, **P < 0.01, ***P < 0.005, Mann-Whitney U test. (B, C, E, F) Data are representative of three independent experiments.

The role of IL-1Ra on Tfr cells was investigated in IL-1Ra^-/-^ mice in the absence of a direct means of eliminating IL-1Ra only in Tfr cells (Fig. 3D-F). At steady state, IL-1Ra deletion did not affect the proportion of Tfh and Tfr cells (Fig. 3D). In addition, circulating IgG were not affected, whereas GCB cells were significantly decreased (Fig. 3D). Four weeks after OVA-immunization, and similarly to Foxp3^creYFP^IL-1R2^lox^ mice, we observed a significant decrease in OVA-specific IgG production, whereas the proportion of GCBs was not significantly altered (Fig.3E, F).

The significant reduction in OVA-specific IgG production despite preserved levels of GCBs reported four weeks after immunization (Fig. 3D) led us to study more deeply the maturation of B cells in response to an antigen challenge. We thus performed deep immunophenotyping of B cells from Foxp3^creYFP^IL-1R2^lox^, Foxp3^GFP^IL-1Ra^-/-^ and control mice at D3, D6 or D9 after immunization using spectral cytometry including CXCR5, CD45R, IgD, CD38, CD95, GL7, IA-IE markers. Multidimensional scaling representations constructed based on the abundance of B cell clusters showed that timepoints were distinguishable in control mice (Fig. 3G and 3H), but overlapped in both Foxp3^creYFP^IL-1R2^lox^ (Fig. 3G) and Foxp3^GFP^IL-1Ra^-/-^ mice (Fig. 3H), suggesting defective maturation of the B cell response between D3 and D9. Overall, our data indicate that IL-1R2 and IL-1Ra on Tfr cells are critical for the production of antigen-specific antibodies. In sum, mice lacking IL-1R2 on Tfr cells have (i) Tfrs showing a more activated phenotype and (ii) decreased antibody production after immunization. Mice lacking IL-1Ra ubiquitously (Arend, 1991) have similar changes in Tfr activation and antibody responses.

### IL-1R2 and IL-1Ra inhibitory receptors prevent IL-1β-mediated activation of Tfr cells

To go further into the characterization of IL-1R2 and IL-1Ra in Tfr cell functions, we developed functional assays where Tfr cells were cultured in the presence of IL-1β. Compared with control Tfr cells, Tfr cells derived from Foxp3^creYFP^IL-1R2^lox^ or Foxp3^GFP^IL-1Ra^-/-^ mice showed increased expression of GITR without IL-1β stimulation (Fig. 4A, B). Upon IL-1β stimulation, expression of GITR was even more enhanced in Tfr from Foxp3^creYFP^IL-1R2^lox^ mice, whereas it remained stable in Tfr from Foxp3^GFP^IL-1Ra^-/-^ mice. Of note, in the absence of IL-1β, IL-1Ra-deleted Tfr cells produced more IL-10 than control Tfr cells (Fig. 4B), whereas production of IL-10 by Tfr from Foxp3^creYFP^IL-1R2^lox^ mice was equivalent to controls (Fig. 4A). IL-10 production was enhanced by the addition of IL-1β in control Tfr cells, whereas it did not affect IL-10 production by Tfr from Foxp3^creYFP^IL-1R2^lox^ and Foxp3^GFP^IL-1Ra^-/-^ mice.

**Fig. 4.**
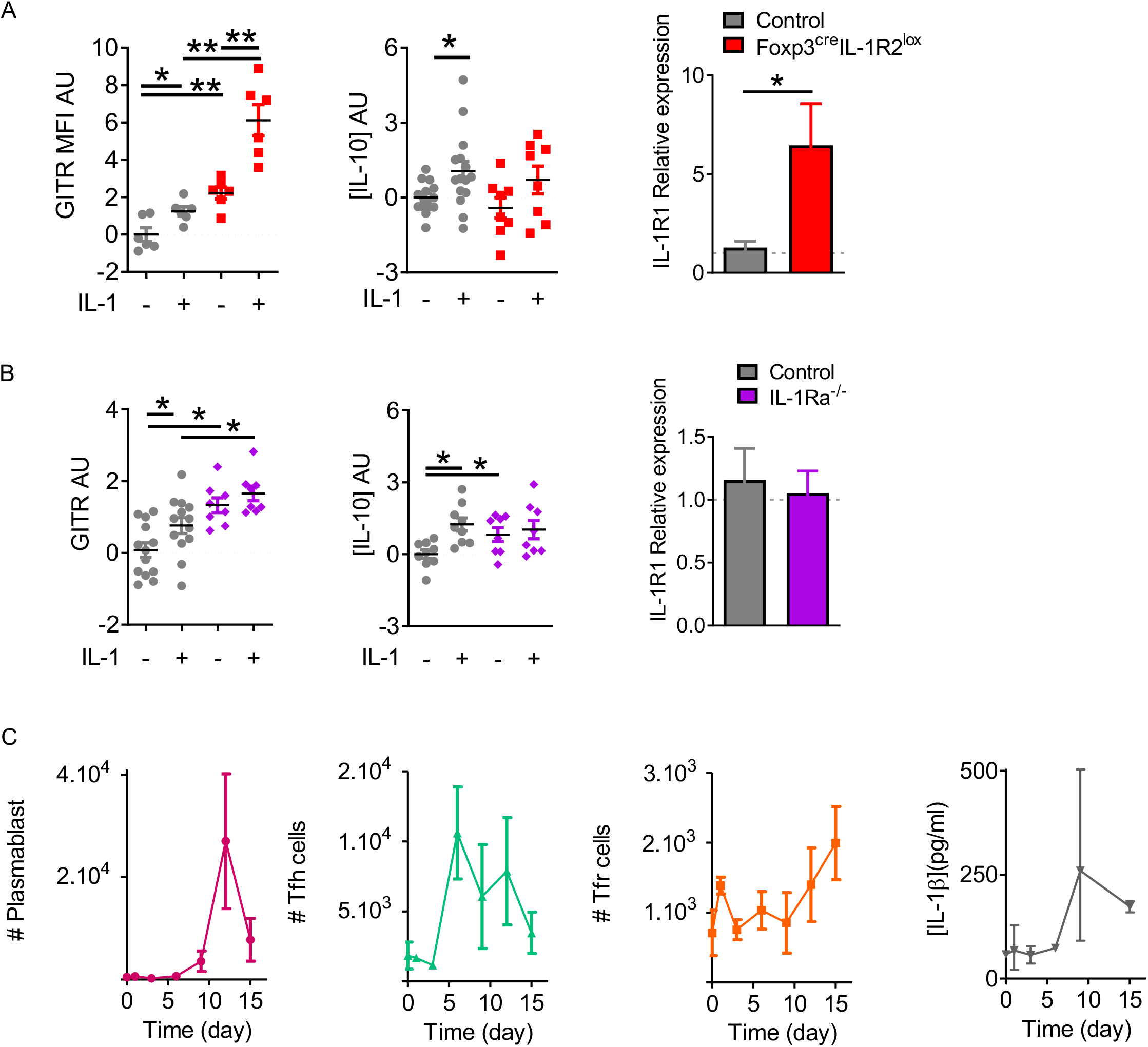
IL-1R2 and IL-1Ra inhibitory receptors prevent IL-1β-mediated activation of Tfr cells. Quantification of GITR expression, IL-10 production and IL-1R1 expression by Tfr cells from (A) Foxp3^creYFP^IL-1R2^lox^ (red square) and Foxp3^creYFP^ (B) Foxp3^GFP^IL-1Ra^-/-^ mice (purple diamond) and Foxp3^GFP^ mice (B) (grey circle), activated with or without IL-1. GITR MFI and IL-10 levels are expressed in arbitrary units that correspond to data centered and reduced relative to Tfr from respective control mice cultivated without IL-1. IL-1R1 expression is evaluated as the relative expression corresponding to the IL-1R1 expression level of the Tfr cell population compared to its relative level in control mice. (C) Kinetics of plasmablasts (pink), Tfh cells (green), Tfr cells (orange) and IL-1β production (grey) in draining lymph nodes from C57BL/6 mice at D0, D3, D6, D9, D12 and D15 after subcutaneous Alum-OVA immunization. Data represent the median of 3 mice ± SEM.*P < 0.05, **P < 0.01, ***P < 0.005, Mann-Whitney U test for unpaired data and Wilcoxon paired test for paired data. (A, B) Data are representative of three independent experiments.

IL-1R2 and IL-1Ra have both extracellular and intracellular actions (Arend, 1991). The low expression of IL-1R1 by T cells makes its detection by flow cytometry difficult, so we quantified IL-1R1 expression by RT-PCR. Noteworthily, IL-1R2 but not IL-1Ra deletion on Tfr cells enhanced their expression of IL-1R1, suggesting a feedback loop of regulation in between IL-1R2 and IL-1R1 (Fig.4 A and B).

Finally, we assessed the dynamics of IL-1β concentrations in the draining lymph nodes of OVA-immunized mice at D0, D3, D6, D9, D12, or D15. IL-1β was detected as early as day 0, but its concentration peaked at D9 (Fig. 4C). This increase preceded the proliferation of Tfr, which appeared after D9 (Fig. 4C). Based on these results, we propose that IL-1 inhibitory receptors are expressed by Tfr to prevent their early own activation by IL-1, which is present at low levels early after immunization. It thus appears that, during evolution, the effects of IL-1 on the GC reaction induced a mechanism to block IL-1R1-mediated activation. This blockage was selected rather than a loss of its expression, highlighting the tinkering nature of evolution (Garlanda et al., 2013; Zheng et al., 2013; Tahtinen et al., 2022).

### Tfr IL-1R expression regulates auto-antibody response in experimental lupus

We then evaluated whether this control of humoral immunity could also affect the autoimmune humoral response in the model of pristane-induced lupus. In this model, the injection of pristane induces the development of anti-DNA autoantibodies in BALB/c mice and anti-RNP antibodies in C57BL/6 mice (Jacob, 1977) and mimics lupus renal damage with the presence of glomerular immune complex deposits at 6 months (Reeves et al., 2009; Simpson et al., 2010). Remarkably, while both Foxp3^creYFP^IL-1R1^lox^ mice and control mice developed anti-RNP antibodies (Fig. 5A), Foxp3^creYFP^IL-1R2^lox^ and IL-1Ra^-/-^ mice did not (Fig 5A).

**Fig. 5.**
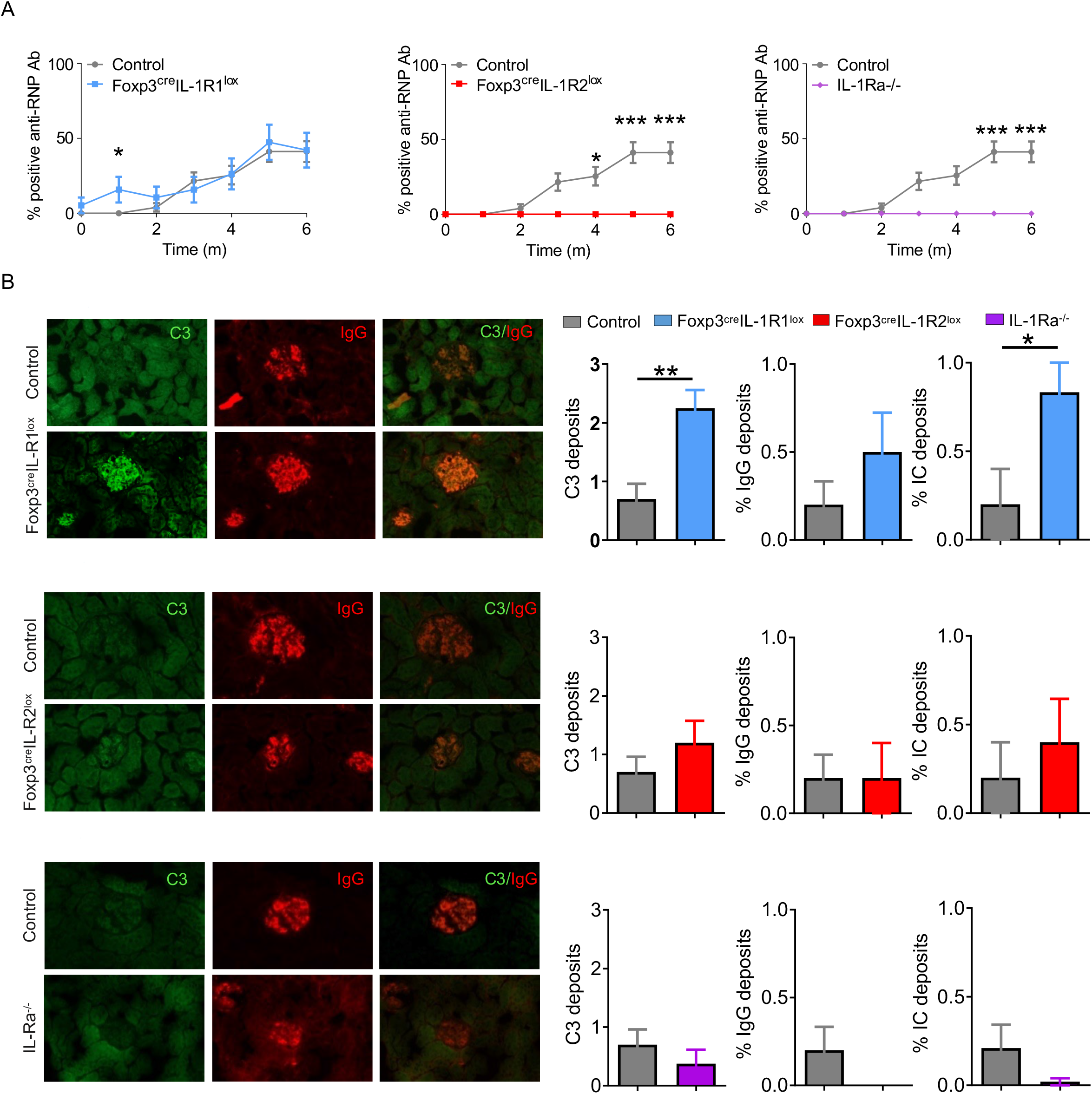
Deletion of IL-1R1 on Tfr cells does not alter the risk of autoimmunity, but deletion of IL-1R2 and IL-1Ra on Tfr cells prevents autoimmune disease. (A) Proportion of mice with positive anti-RNP Ab after pristane immunization in Foxp3^creYFP^IL-1R1^lox^ (blue, n= 6), Foxp3^creYFP^IL-1R2^lox^ (red, n= 6), IL-1Ra^-/-^ (purple, n= 6) and control (grey, Foxp3^YFP^, Foxp3^YFP^ or and Foxp3^GFP^ respectively, n= 6) mice. (B) Deposition of C3 (green) and IgG (red) in glomeruli of kidney tissues from Foxp3^creYFP^IL-1R1^lox^ (blue, n=6), Foxp3^creYFP^IL-1R2^lox^ (red, n=5), IL-1Ra^-/-^ (purple, n=5) and control (grey, Foxp3^creYFP^and Foxp3^GFP^, n=10) mice exposed to pristane. Intensity of C3 deposition is evaluated using a scale ranging from 0: absent, 1: weak, 2: moderate to 3: intense and IgG deposition is defined as an IgG immunofluorescence score greater than 2. The proportion of mice with immune complex deposition is defined as IgG and C3 immunofluorescence score greater than 2 in Foxp3^creYFP^IL-1R1^lox^ (blue), Foxp3^creYFP^IL-1R2^lox^ (red), IL-1Ra^-/-^ (purple) and control (grey, Foxp3^YFP^ and Foxp3^GFP^) mice after 6 months of exposure to pristane. *P < 0.05, **P <0.01, ***P < 0.005, Mann-Whitney U test. (A, B) Data are representative of three independent experiments.

We then analyzed glomerular damage at 6 months after disease induction by evaluating immune complex deposition. Foxp3^creYFP^IL-1R1^lox^ mice showed more glomerular C3 and immune complex depositions than control mice, while the proportion of glomeruli affected by IgG deposition was not significantly increased (Fig. 5B). Foxp3^creYFP^IL-1R2^lox^ mice showed similar rates of C3, IgG, or immune complex deposition as controls (Fig. 5B), despite the absence of anti-RNP production. Similarly, IL-1Ra^-/-^ mice showed a non-significant decrease of immune complex deposition. Altogether, results indicate that deletion of IL-1R2 and IL-1Ra on Tfr abolishes the production of anti-RNP antibodies, which does not translate into significant glomerular immune complex deposits, whereas IL-R1 deletion on Tfr enhances lupus renal damage, confirming defective regulation of Ab production in Foxp3^creYFP^IL-1R1^lox^ mice. In C57/BL6 mice, pristane induces a pauci-immune clinical form with low anti-RNP production (and very low anti-DNA antibodies), but with potentially severe renal damage (Huang et al., 2017). Indeed, while the level of autoantibodies produced by Foxp3^creYFP^IL-1R1^lox^ mice was only slightly increased one month after disease induction, we found more severe renal damage compared to control mice. Conversely, although the level of autoantibodies was completely abolished in Foxp3^creYFP^IL-1R2^lox^ and IL-1Ra^-/-^ mice, renal damage was not significantly reduced. Nevertheless, our results show a role of IL-1R expression in both disease severity and autoantibody production.

Taken together, our data indicate a fine regulation of IL-1β within the GC: IL-1 can activate both Tfh and Tfr cells, with important implications for the regulation of humoral immunity. Indeed, due to the concomitant expression of IL-1R2 and IL-1Ra on Tfr cells, the latter can only be activated at higher concentrations of IL-1. Taking into account the kinetics of IL-1 production, which progressively increases in lymph nodes after immunization, we propose that IL-1 activation of Tfr should occur after that of Tfh, ensuring the regulation of antibody production.

## Material and methods

### Study design

For flow cytometry assays, the sample sizes were of at least three individuals per experiment. For enzyme-linked immunosorbent assay (ELISA), sample sizes were of at least three per condition. Sample size was determined on the basis of experimental feasibility and statistical significance. The experiments were not randomized. The investigators were not blinded to the allocation during experiments and analyses.

### Mice

B6.129S-IL1rntm1Dih/J (JAX stock #004754), B6.Cg-Tg(Cd4-cre)1Cwi/Bflu/J (JAX stock #022071), B6.129(Cg)-Il1r1tm1.1Rbl/J (JAX stock #028398), B6.129(Cg)-Foxp3tm4(YFP/icre)Ayr/J (JAX stock #016959) mice were provided by the Jackson Laboratory. C57BL/6N-Atm1BrdIl1r2tm1a(EUCOMM)Wtsi/WtsiPh (MGI:4842437) were provided by Institute of Molecular Genetics of the ASCR. Confirmed genotyping was performed in accordance with the supplier recommendations. All transgenic mice were in a C57BL/6J background. C57BL/6 Foxp3-GFP mice expressing GFP under the control of the promoter of Foxp3 gene were provided by B. Malissen of the Centre d’Immunologie de Marseille-Luminy (France). To generate CD4^cre^IL-1R1^lox^ mice, Il1r1tm1.1Rbl/J mice were mated with B6.Cg-Tg(Cd4-cre)1Cwi/Bflu/J mice. To generate FoxP3^creYFP^IL-1R1^lox^ mice Il1r1tm1.1Rbl/J mice were mated with B6.129(Cg)-Foxp3^tm4(YFP/icre)Ayr^/J mice. To generate Foxp3^GFP^CD4^cre^IL-1R1^lox^ mice, allowing to distinguish Tfh and Tfr cells, CD4^cre^IL-1R1^lox^ mice were crossed with Foxp3^GFP^ mice. To generate Foxp3^GFP^IL-1Ra^-/-^ mice, B6.129S-IL1rntm1Dih/J mice were crossed with Foxp3^GFP^ mice. To generate FoxP3^creYFP^IL-1R2^lox^ mice, C57BL/6N-A^tm1BrdI^l1r2^tm1a(EUCOMM)Wtsi/Wtsi^ mice were mated with B6.129(Cg)-Foxp3^tm4(YFP/icre)Ayr^/J mice. All animals were maintained at the University Pierre and Marie Curie (UPMC) Centre d’Expérimentation Fonctionnelle animal facility (Paris, France) under specific pathogen–free conditions in agreement with current European legislation on animal care, housing, and scientific experimentation (agreement number A751315). All procedures were approved by the local animal ethics committee.

### Immunization models

#### OVA Immunization models

Mice were either immunized with intraperitoneal injection three times (D0, D2, D4) and sacrificed at D8 for Tfr /Tfh cell transcriptomic and in vitro studies or immunized twice (D0 and D14). Intraperitoneal injection was performed with 100 ug of OVA (OVA A5503, Sigma-Aldrich) mixed with 500 μg of aluminum hydroxide (Alum) gel (AlH303, Sigma). Mice were also immunized with OVA-Alum subcutaneously (s.c), in the flank one time and sacrificed at D0, D3, D6, D9, D12, D15 for deep immunophenotyping of follicular cells or to measure the level of IL-1 from inguinal draining lymph.

#### Pristane Immunization model

Lupus was induced by intraperitoneal injection with 100 μg of Pristane at D0. Mice were euthanized at 6 months (M6) for the quantification of immune complex deposition in the kidney.

### Cell sorting

Splenocytes from immunized mice were stained with Ter-119–biotin, CD8-biotin, CD11c-biotin and B220-biotin antibodies for 20 min at 4°C and labeled with anti-biotin magnetic beads (Miltenyi Biotec) for 15 min at 4°C. Biotinylated cells were depleted on an autoMACS separator (Miltenyi Biotec), following the manufacturer’s procedure. Enriched T cells were stained as described in the “Flow cytometry analysis of human cells” section, and the following subsets were sorted on BD FACSAria II (BD Biosciences), with a purity of >98%: CD4^+^CD8^-^CXCR5^hi^PD1^hi^Foxp3^-^ Tfh cells, CD4^+^CD8^-^CXCR5^hi^PD^hi^Foxp3^+^ Tfr cells. Inguinal lymph node were comminuted mechanically and filtered through a cell strainer (35 μm, StemCell). The diluent was stored at −80°C to perform IL-1β ELISA. The cell suspension was transferred into a 5 mL tube (Falcon) containing PBS, washed twice with PBS medium and stained.

### Analysis of lupus kidney involvement and research of immune complex deposition

One kidney/mouse was embedded in OCT Tissue-Tek compound (CellPath), snap-frozen, and stored at −80°C until use. Kidney were frozen with optimal cutting temperature (OCT) compound, cut into 7 μm sections and fixed with acetone. Sections will then be stained with FITC-conjugated anti-C3 and Alexa Fluor 647-conjugated anti-mouse IgG (Abcam, Cambridge, UK) antibodies. DNA was visualized using DAPI (Sigma-Aldrich). Quantification of fluorescence will be performed at ×20 magnification in one random microscopic field of each kidney, followed by binary analysis using Image J 1.49v software. Scores on a scale of 0–3 were estimated for both C3 and IgG deposition, based on the fluorescence intensity (0 = no deposits, 1 = low, 2 = moderate, 3 = high). An intensity threshold of 2 was considered positive for IgG or D3 deposits. Immune complex deposits positivity was assessed by a score above 2 for IgG and C3.

### Flow cytometry analysis

Fresh total cells from spleens were isolated in PBS1×–3% fetal bovine serum (FBS) and stained for 20 min at 4°C with the following monoclonal antibodies at predetermined optimal dilutions: CD121b-BV421, CD19-PeCF594, CD4-V500, CD8a-AF700, Bcl6-APC, CXCR5-Biotin, GL7-e450, CD95 PE, Foxp3-AF488, PD-1–PE (BD Biosciences) or PE-Texas Red (PETR), streptavidin-APC or streptavidin–APC-Cy7 (BD Biosciences), GITR PETR (Miltenyi). CXCR5 staining was performed using biotinylated anti-CXCR5 for 30 min at 20°C followed by APC- or APC-Cy7–labeled streptavidin at 4°C. Intracellular detection of Foxp3 was performed on fixed and permeabilized cells using appropriate buffer (eBioscience), following the manufacturer’s recommendations. Stained cells were run on CytoFLEX S cytometer (Beckman - Coulter) and analyzed using FlowJo software (TreeStar Inc.). Dead cells were excluded by forward/side scatter gating.

### Full Spectrum Flow Cytometry profiling

Fluorophores-conjugated antibodies are listed in Table 1. Two million cells per sample were stained. Cell viability was assessed by Viability UV T (Thermofisher). Cells were washed with PBS. After washing, anti-chemokine antibody (CXCR5) and brilliant stain buffer (used in complement of Brilliant fluorescent polymer dyes) were added in PBS 1 × at room temperature for 20minutes. Other membrane markers were added next (CD44, CD69, ICOS, CD40L, OX40, Fas, GL7, CTLA4, GITR, IgD, IgG, IgM, CD38, B220, CD19) and lymph node cells were incubated for 30 additional minutes at room temperature. After washing, the cells were washed and then resuspended in a residual volume of 100uL and acquired on an AURORA FSFC (Cytek Biosciences, Fremont, CA). Supervised analyses, conducted using FlowJo software, were used to study the phenotype and function of some specific cell populations.

**Table 1.** Full spectrum flow cytometry antibody panel.

| Label | Target | Clone | Providers |
| --- | --- | --- | --- |
| BUV 737 | CXCR5 | 2G8 (RUO) | Becton Dickinson |
| AF647 | CD19 | 6D5 | Biolegend |
| APC-Fire810 | CD45R | RA3-6B2 | Biolegend |
| APC-R700 | CD4 | RM4-4 | Biolegend |
| SparkBlue 550 | CD8 | 53-6.7 | Biolegend |
| BB700 | CD11b | M1/70 (RUO) | Becton Dickinson |
| BV480 | CD28 | 37.51 (RUO) | Becton Dickinson |
| BUV615 | CD40L | MR1 (RUO) | Becton Dickinson |
| BUV496 | CD69 | H1.2F3 (RUO) | Becton Dickinson |
| BUV661 | OX40 | ACT35 (RUO) | Becton Dickinson |
| APC | CD152/CTLA4 | UC10-4F10-11 (RUO) | Becton Dickinson |
| PerCP-Cy5.5 | GL7 | GL7 | Biolegend |
| BUV615 | IgD | 751474 (RUO) | Becton Dickinson |
| BV650 | CD25 | PC61 (RUO) | Becton Dickinson |
| PerCP | CD3 | 145-2C11 | Biolegend |
| BUV395 | CD38 | 90/CD38 | Becton Dickinson |
| BV570 | CD44 | IM7 | Biolegend |
| PE-CF594 | CD95 | Jo2 (RUO) | Becton Dickinson |
| BV711 | GITR | DTA-1 | Becton Dickinson |
| PC5 | ICOS | 15F9 | Biolegend |
| APC | IgM | RMM-1 | Biolegend |
| BV786 | PD1 | RMP1-30 (RUO) | Becton Dickinson |
| BUV805 | IA-IE | M5/114.15.2 | Becton Dickinson |
| BUV563 | IgG1 | A85-1 (RUO) | Becton Dickinson |
| UBUV420 | Viability UV T |  | Thermo Fisher |
| PE | CD121a | 35F5 (RUO) | Becton Dickinson |
| BV421 | CD121b | 4E2 (RUO) | Becton Dickinson |

For unsupervised analysis used in the MDS representation, batch corrections of expression values for all markers and cell clusters analyses were performed using Imubac R package. We generated 40 cell clusters based on the expression of cell markers. We manually determined the identity of these clusters and selected B cell clusters. The MDS representations were generated based on the abundances of B-associated cell clusters in each mouse of subsequent analyses.

from their locations in the UMAP plot and the heatmap summarizing the median expression levels of all markers for each cluster.

### Gene expression analysis based on RT-PCR

Sorted cells were washed in PBS1× and stored in RNAqueous kit lysis buffer (Ambion Inc./Life Technologies) at −80°C. Total RNA was extracted according to the manufacturer’s instructions, and quality was assessed on a bioanalyzer using the Pico RNA Reagent Kit (Agilent Technologies). For quantitative PCR, we analysed 25 ng of cDNA for expression of a range of genes using the GeneAmp 5700 Sequence Detection System (Applied Biosystems). The samples were normalized to the GADPH protein as control mRNA, by change in cycling threshold (ΔCT) method and calculated based on 2-ΔΔCT.

### In vitro assay

Tfr cells were sorted from D-8 OVA-immunized mice, following a previously described “Immunization” protocol. For all the conditions, 1 × 10^4^ Tfr cells were cultured for 12 hours in 96-well plates (Nunc) in complete RPMI 1640 (Thermo Scientific) and three CD3/CD28 beads for one T cell (Dynabeads Mouse T-Activator, Thermo Scientific). We then added or not recombinant mouse 1 μg of IL-1β (Milteniy Biotec) to Tfr cell cultures. IL-10 production by Tfr cells was measured by ELISA (eBioscience) in supernatants of cultured Tfr cells. Expression of activation marker (GITR) was analyzed by flow cytometry.

### ELISA

The concentrations of mouse IL-10 (eBiosciences), IL-1Ra (eBiosciences), IL-1β (eBiosciences) in the culture supernatant, lymph node lavage fluid and serum were detected using ELISA kits. The serum levels of anti-OVA and anti-RNP antibodies were detected with in-house ELISA. Briefly, 5 ng/ml OVA proteins were coated on 96-well high-binding EIA/RIA plates overnight. After blocking, mouse serum that had been diluted at different concentration (1:4000, 1:12000, 1:36000, 1:108000, 1:324000, 1:972000) was added to the plates and incubated for 2 h; then HRP-conjugated anti-mouse IgG (Southern Biotech) was added at a 1 : 2500 dilution followed by Streptavidin revelation. The anti-RNP antibodies were revealed with similar method: 1,33 μg/ml RNP proteins were coated on 96-well high-binding EIA/RIA plates overnight. After blocking, mouse serum that had been diluted at different concentration (1:100, 1:300, 1:900, 1:2700) was added to the plates and incubated for 2 h; then HRP-conjugated anti-mouse IgG (Southern Biotech) was added at a 1: 2500 dilution followed by Streptavidin revelation. For total IgG levels in serum, Maxisorp (Nunc) plates were coated with anti-mouse Ig (ref: 0107-01, SouthernBiotech) on 96-well high-binding EIA/RIA plates overnight. After blocking, mouse serum that had been diluted at different concentration was added to the plates and incubated for 2 h; then HRP-conjugated anti-mouse IgG (ref: 1030-01, Southern Biotech) was added at a 1: 2500 dilution followed by Streptavidin revelation.

### Statistical analysis

Flow cytometry, cytokine production, and gene expression data were analyzed by nonparametric Mann-Whitney U test for unpaired data and paired Wilcoxon for paired data on GraphPad Prism v5 [P values, such as P > 0.05 (not significant), *P < 0.05, **P < 0.01, ***P < 0.001, and ****P < 0.0001, are indicated in the figures]. Spectral cytometry data were analysed after batch correction with Imubac package.

## List of Supplementary Materials

Fig. S1. Phenotype of KO mice

## Acknowledgements

We thank Olivier Jaen and Flora **Guillot (Cytek Biosciences);** Bénédicte Hoareau (UMS 37 PASS); Gwendolyn Marguerit, Encarnita Mariotti-Ferrandiz, Gabriel Pires and Thomas Vasquez (i3 lab); Doriane Foret, Flora Issert, Kim Nguyen, Olivier Bregerie (UMS 28). Funding: AB was supported by the Fondation pour la Recherche Médicale and PE by a grant from the Swiss National Science Foundation and the Novartis Research Foundation.

## Supplementary Materials

**Fig. S1.**

(A) Relative expression of IL-1R1 in Tfol (CD4^+^CD19^-^PD1^+^CXCR5^+^), Tfh (CD4^+^Foxp3^-^CD19^-^PD1^+^CXCR5^+^) and Tfr (CD4^+^Foxp3^+^CD19^-^PD1^+^CXCR5^+^) cells from Foxp3^GFP^CD4^cre^IL-1R1^lox^ (green) and Foxp3^GFP^(grey) mice by RT-PCR. IL-1R1 expression is evaluated as the relative expression corresponding to the IL-1R1 expression level of a cell population compared to its relative level in control mice. (B) Relative expression of IL-1R1 in Tfh and Tfr cells of Foxp3^creYFP^IL-1R1^lox^ (blue) and Foxp3^YFP^ (grey) mice by RT-PCR. IL-1R1 expression is assessed as previously mentioned. (C) Quantification of IL-1R2 expression on Tfr, Tfh, Treg (gated as CD4^+^Foxp3^+^CD19^-^PD1^-^CXCR5^-^) and Teff (gated as CD4^+^Foxp3^-^CD19^-^PD1^-^CXCR5^-^) cells from Foxp3^GFP^ control mice. (D) Quantification of IL-1R2 expression on Tfr, Treg and Breg cells (CD4^-^CD19^+^Foxp3^+^) from Foxp3^creYFP^IL-1R2^lox^ and Foxp3^creYFP^ mice. (E) Blood IL-1Ra expression in IL-1Ra^-/-^ (purple) and Foxp3^GFP^(grey) mice after PBS or OVA-Alum immunization. *P < 0.05, **P < 0.01, ***P < 0.005, Mann-Whitney U test.

