## Supplementary figures and images for "Interleukin-1 regulates follicular T cells during the germinal center reaction"

### Supplemental Figure 1

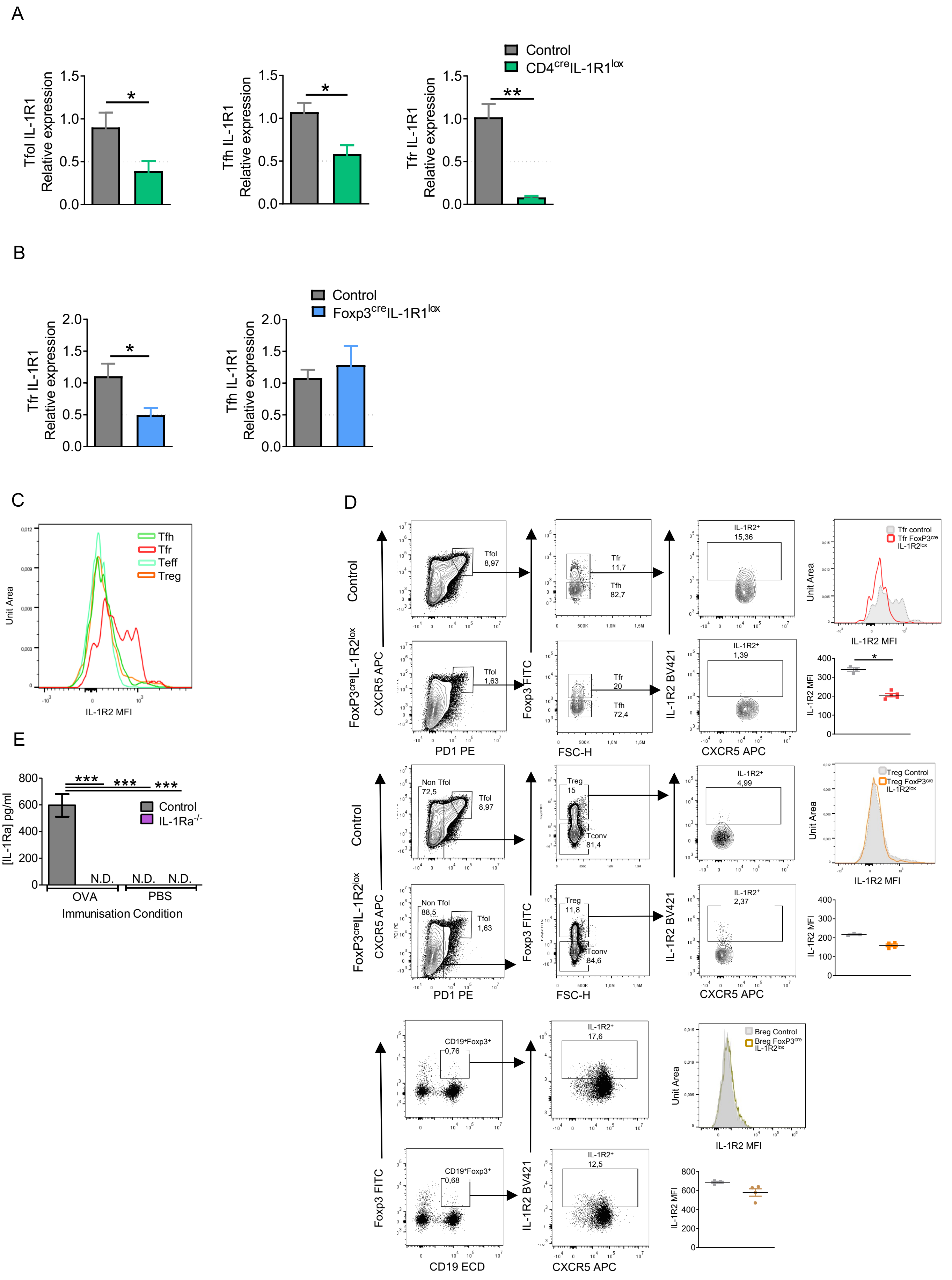
